# Novel effector genes revealed by the genomic analysis of the phytopathogenic fungus *Fusarium oxysporum* f. sp. *physali* (*Foph*) that infects cape gooseberry plants

**DOI:** 10.1101/2020.08.10.235309

**Authors:** Jaime Simbaqueba, Edwin A. Rodriguez, Diana Burbano-David, Carolina Gonzalez, Alejandro Caro-Quintero

## Abstract

The vascular wilt disease caused by the fungus *Fusarium oxysporum* f. sp. *physali* (*Foph*) is one of the most limiting factors for the production and export of cape gooseberry (*Physalis peruviana*) in Colombia. A previous study of the transcriptomic profile of a highly virulent strain of *F. oxysporum* in cape gooseberry plants, from a collection of 136 fungal isolates obtained from wilted cape gooseberry plants, revealed the presence of secreted in the xylem (SIX) effector genes, known to be involved in the pathogenicity of other *F. oxysporum formae speciales* (ff. spp.). This pathogenic strain was named *Foph*, due to its specificity for cape gooseberry hosts. Here, we sequenced the genome of *Foph*, using the Illumina MiSeq platform. We analyzed the assembled genome, focusing on the confirmation of the presence of homologues of SIX effectors and the identification of novel candidates of effector genes unique of *Foph.* By comparative and phylogenomic analyses based on single-copy orthologues, we identified that *Foph* is closely related to *F. oxysporum* ff. spp., associated with solanaceous hosts. We confirmed the presence of highly identical homologous genomic regions between *Foph* and *Fol*, that contain effector genes and identified seven new effector gene candidates, specific to *Foph* strains. We also conducted a molecular characterization of a panel of 29 *F. oxysporum* additional stains associated to cape gooseberry crops isolated from different regions of Colombia. These results suggest the polyphyletic origin of *Foph* and the putative independent acquisition of new candidate effectors in different clades of related strains. The novel effector candidates identified by sequencing and analyzing the genome of *Foph*, represent new sources involved in the interaction between *Foph* and cape gooseberry. These resources could be implemented to develop appropriate management strategies of the wilt disease caused by *Foph* in the cape gooseberry crop.

## Introduction

*Fusarium oxysporum* is a cosmopolitan ascomycete fungus that commonly inhabits agricultural soils. Rather than a single species, it is a species complex of non-pathogenic, plant pathogenic, and human pathogenic strains, termed the *Fusarium oxysporum* species complex (FOSC) (Di Pietro et al., 2003; Michielse and Rep 2009; O’Donnell et al. 2009; Ma et al., 2013, Ma, 2014). Several hundred different members of the FOSC are able to penetrate plant roots, colonise xylem vessels and produce vascular wilt diseases in a broad range of host plants, including economically important crops such as banana, cotton, date palm, onion, brassicas, cucurbits, legumes and solanaceous species, such as tomato, eggplant, chilli and cape gooseberry, but not grasses (Michielse and Rep, 2009). However, individual pathogenic isolates of *Fusarium oxysporum* are highly host specific and have therefore been classified into different *formae speciales* (ff. spp.) according to the host they infect e.g. strains that infect banana cannot infect tomato plants and vice versa (Lievens et al., 2008; Michielse and Rep 2009; Ma, 2014). *F. oxysporum* has no known sexual stage and the mechanism for species diversification has been associated with the parasexual cycle through heterokaryon formation, which enables a mitotic genetic exchange between different nuclei (Glass et al., 2000; Di Pietro et al., 2003).

Comparative genomics of phytopathogens in the genus *Fusarium* (i.e. *F. graminearum, F. verticillioides* and *F. oxysporum* f. sp. *lycopersici* [*Fol*]), revealed the presence of lineage specific (LS) chromosomes and chromosomal regions in *Fol* that were rich in repetitive elements and contained genes encoding known or putative effector proteins (Ma et al., 2010). Among them, 14 genes were identified that encode small proteins secreted into the xylem sap of tomato plants infected with *Fol* (called SIX proteins) (Houterman et al., 2007; Schmidt et al., 2013). Three of these *SIX* genes are avirulence genes *(Avr)*, with resistance *(R*) gene counterparts identified in tomato (Simons et al., 1998; Rep et al., 2004; Houterman et al., 2008, 2009; Catanzariti et al., 2015, 2017).

Small proteins secreted by a broad range of plant pathogens, including bacteria, fungi, oomycetes and nematodes, that interfere with the cellular structure and function of their hosts are known as effector proteins (Kamoun, 2006, 2007; Hogenhout et al., 2009). The low level of homology among fungal effectors makes it difficult to identify common features that allow their classification as a group or protein family (Stergiopoulos and de Wit, 2009; Lo Presti et al., 2015; Guillen et al., 2015). Nevertheless, many fungal effectors have been identified based on the presence of a signal peptide sequence for secretion, small size of around 300 amino acids or less, and the fact that they are often cysteine rich (Sperschneider et al., 2015). A large-scale search for putative effector genes in 59 strains of various ff. spp., resulted in a set of 104 candidate effectors including the 14 secreted in the xylem (SIX) genes, identified in *Fol* (Ma et al., 2010; Schmidt et al., 2013; van Dam et al., 2016). From this candidate effector repertoire, strains were classified according to the putative effector sequences they shared. Interestingly, all the cucurbit-infecting ff. spp. *(melonis, niveum, cucumerinum* and *radicis-cucumerinum*), were grouped together in a separate supercluster, sharing an overlapping set of putative effectors and possibly conferring the ability to those ff. spp. to infect cucurbit host species (van Dam et al., 2016). This supercluster includes a substantial overlap with *SIX1, SIX6, SIX8, SIX9, SIX11 and SIX13* and largely excluded *SIX2, SIX3, SIX4, SIX5, SIX7, SIX10, SIX12* and *SIX14.* Homologues of *Fol* SIX genes have been identified in alliaceous, legumes, musaceous, solanaceous and narcissus infecting ff. spp. of *F. oxysporum* (Taylor et al., 2016 and 2019; Williams et al., 2016; Czislowski et al., 2018; Simbaqueba et al., 2018).

Cape gooseberry (*Physalis peruviana*) from the Solanaceae family, is a tropical native fruit of South America found typically growing in the Andes. In Colombia, over the last three decades, the cape gooseberry has been transformed from a wild and under-utilized species to an important exotic fruit for national and international markets and represents one of the most exported fruit for Colombia (Simbaqueba et al., 2011; Moreno-Velandia et al., 2019). The cape gooseberry is also appreciated by its nutritional and medicinal properties (Yen et al., 2010; Ramadan 2011, 2015; El-Gengaihi et al., 2013). However, despite its significant value, cape gooseberry production has been limited due to the lack of known cultivars and the absence of adequate phytosanitary measures. One of the most important disease problems in cape gooseberry is the vascular wilt disease caused by *Fusarium oxysporum*. This disease was first described in 2005 and has become one of the limiting factors for cape gooseberry production and export (Moreno-Velandia et al., 2018). Field observations indicated typical symptoms of a vascular wilt disease with an incidence ranging from 10 to 50% with losses in production of 90% approximately (unofficially reported), in the Cundinamarca central region of Colombia. Consequently, producers moved to other places in the same region, spreading contaminated plant material and seeds (Barrero et al., 2012).

From 2012 to 2015, a total of 136 fungal isolates were obtained from cape gooseberry plants showing wilting disease symptoms, collected from different locations of the central Andean Region of Colombia. The fungal isolates were described as *F. oxysporum*, using Koch postulates and molecular markers for intergenic spacers (IGS) and the Translation Elongation Factor 1 Alpha gene (EF1α) of *F. oxysporum* (AGROSAVIA, Unpublished results). From the *F. oxysporum* strains, one (named MAP5), was found to be highly virulent on a commercial variety of cape gooseberry and different accessions from National Germplasm Bank and different collections (Enciso-Rodriguez et al., 2013; Osorio-Guarin et al., 2016). Further RNAseq analysis was performed to study differential gene expression comparing susceptible and resistant cape gooseberry plants inoculated with MAP5 (AGROSAVIA, Unpublished results). This RNAseq data was used in comparative transcriptomics, identifying eight homologues of effector genes between *Fol* and MAP5. Thus, describing a newly forma specialis of *Fusarium oxysporum* that affect cape gooseberry plants, designated as *F. oxysporum* f. sp. *physali (Foph*) (Simbaqueba et al., 2018).

In this study, we sequenced the genome of *Foph* and performed comparative genomics using the resulted genome assembly to infer the phylogenetic relationship of *Foph* within the *F. oxysporum* clade. This result showed the polyphyletic origin of *Foph* and the closer relationship with ff. spp. related to *Solanaceous* hosts. We also identified putative LS genomic regions in *Foph* that could be related with pathogenicity and host specificity, as they contain the homologous effectors previously reported and eight new effector candidates identified in this study. We mapped the *Foph* RNAseq dataset previously reported against the candidate effectors and identified that these novel effectors are expressed during host infection. These results indicate that the new candidate effectors, could have a putative role in virulence. Additionally, we tested the presence of the novel effectors by PCR amplification in a panel of 36 *F. oxysporum* isolates (including MAP5), associated to the cape gooseberry crop and identified that the presence of novel candidates was unique to *Foph* related strains, suggesting host specificity towards cape gooseberry plants. Furthermore, we conducted a phylogenetic analysis using the EF1a sequences available for this panel of *F. oxysporum* isolates. This result reflects the polyphyletic origin of *Foph* and suggests the independent acquisition of the candidate effectors in at least two divergent clades of *Foph* related strains.

## Materials and Methods

### DNA extraction

*F. oxysporum* strains were reactivated in PDA media and incubated at 28 °C for eight days or until enough biomass was obtained for DNA extraction. The DNA of MAP5 strain (*Foph*), used for genome sequencing, was obtained using the ZR Fungal / Bacterial DNA kit from Zymo research®, according to the protocol proposed by the manufacturer. The DNA of the remaining *F. oxysporum* isolates used in this study, was extracted from 100 mg of the mycelia, using the cetyltrimethylammonium bromide (CTAB) protocol modified for fungal DNA (Zhang et al., 2010). The quality and DNA concentration using both methodologies were verified in 1% agarose gel using the 1Kb Plus DNA Ladder (Invitrogen®) and also by Nanodrop DNA/RNA Quantification system.

### *Foph* genome sequencing and assembly

Libraries of the virulent strain MAP5 of *Foph*, were generated from purified DNA with the Illumina Nextera XT DNA Sample Preparation Kit (San Diego, California, USA). The resulted libraries were verified in the Bioanalyzer Agilent 2100, using a DNA-HS chip and adjusted to a final concentration of 10 nM. Libraries were then amplified. The sequencing of the libraries was performed using the TruSeq PE Cluster V2 (Illumina, San Diego CA) kit generating 250bp pair-end reads in the Illumina MiSeq platform (San Diego, California, USA) at the Genetics and Antimicrobe Resistance Unit of El Bosque University.

The quality of the reads produced was verified with the software FastQC (Andrews S, 2015), and reads were trimmed using the software Trimmomatic (Bolger et al., 2014), with the following parameters “LEADING:3 TRAILING:3 SLIDINGWINDOW:4:15 MINLEN:45”. Additionally, adaptor sequences and reads less than 25 bp in length were filtered and removed using the scripts fastq_quality_trimmer and fastq_quality_filter of the FASTX-toolkit platform (hannonlab.cshl.edu/fastx_toolkit). A primary *de novo* assembly was performed with the pair-end reads overlapped into contigs, using the software Newbler v 2.0.01.14. (454 Life Sciences), Velvet (Zerbino & Birney, 2008), and SPAdes, v 3.5.0. (Illumina, San Diego CA). The Quality Assessment Tool for Genome Assemblies (QUAST) software (Gurevich et al., 2013), was used to determine the best genome assembly based on the highest N50 parameter.

### Gene prediction and annotation

*Ab initio* gene models for the genome sequence of *Foph*, were predicted using the software Augustus (Stanke & Morgenstern, 2005), using the gene prediction model for *Fol*4287, as species gene model with the following parameters “--strand=both” and “--uniqueGeneId=true”, other parameters were used with the default settings. The resulted transcripts were annotated by combining predictions using the software HMMER 3.0 (Finn et al., 2011), with the PFAM protein database. The functional annotation of the transcripts was performed with the software eggNOGmapper v4.5.1 (Huerta-Cepas et al., 2017). Gene models were corroborated with the *Foph in planta* RNAseq database reported in our previous study.

### Comparative genomics analysis

The comparative genomic analysis was carried out to establish the gene composition similarity and conserved patterns within phylogenetic clusters of 22 genomes of different *F. oxysporum* ff. spp. (including *Foph)*, and the genome sequence of *F. fujikuroi* (Supplementary table S1). To identify these gene clusters, we used the anvi’o software (Eren et al., 2015), following the pangenomic workflow described before (Delmont & Eren, 2018). In brief, this pipeline generates a genome database that stores the DNA and amino acid sequences information of all genomes. Gene clusters were identified by calculating the similarities of each amino acid sequence in every genome against every other amino acid sequence using blastp (Altschul et al., 1990) and finally a hierarchical clustering was performed using the Euclidean distance and Ward clustering algorithm. The distribution of these gene clusters across the genomes was plotted using the anvi’o visualization tool. To reconstruct the phylogenetic relationship of these genomes, the single copy orthologous genes (SCG) were extracted from the pangenome database for all genomes, and a phylogenomic tree was generated using the FastTree 2.1 software (Price, et al., 2010) as a component of the anvi’o pipeline. To root the tree, we used the genome sequence of *F. fujikoroi* as outgroup (Supplementary table S1).

### Identification of effector genes in *Foph*

To validate the presence of homologous effectors (i.e. SIX, Ave1 and FOXM_16303), identified in our previous *Foph* transcriptomic analysis, we carried out two search strategies of the homologue effectors in a database that included the 22 genome sequences of the ff. spp. of *F. oxysporum* used for comparative genomics of *Foph.* The first strategy consisted in a tBlastn search. The hits with an e-value <0.0001 and identity higher than 50%, in the 50% of the length of the sequence query, were selected for further analysis. In the second strategy, a Blastx search was performed to identify all possible putative peptides of the homologous effectors in the *F. oxysporum* genome database. The best hits with an e-value <0.0001 were selected for further analysis (Table 2).

To identify *de novo* candidate effector genes in *Foph*, the secretome and effectorome were predicted from the proteome of *Foph*, using the software SignalP v5.0 (Almagro-Armenteros et al., 2019) and EffectorP v2.0 (Sperschneider et al., 2018), respectively. In order to discard homologous sequences in other *ff. spp.*, The two BLAST search strategies mentioned above were performed using the protein sequences positive for signal peptide and effector structure (i.e. <300 aa in length and cysteine rich) as a query. An additional search of Miniature Impala Transposable Elements *(mimp*) was performed in the UTR of the transcripts predicted of *Foph*, with the regular expression ‘NNCAGT[GA][GA]G[GAT][TGC]GCAA[TAG]AA’, using a customised Perl script as described by Schmidt et al, (2013) and van Dam and Rep, (2017), to determine whether or not the novel candidate could correspond to SIX type genes.

### Molecular characterization of *Foph* isolates and PCR analysis of candidate effectors

A panel of 36 *F. oxysporum* isolates (including the highly virulent *Foph*), derived from a collection of 136 fungal isolates obtained from cape gooseberry crops, were selected based on their ability to cause wilting symptoms on susceptible cape gooseberry plants (Supplementary Table S2). The *EF1a* gene of *Fol* (GenBank XM_018381269), was used as a molecular marker to characterize the *Foph* isolates to species level. EF1a sequences for seven out of 36 isolates (including MAP5), were obtained from the GenBank (Supplementary Table S2). For the remaining 29 isolates, a fragment of the EF1a gene was amplified and sequenced using the primers reported by Imazaki et al, (2015). PCR reactions were conducted with Taq DNA Polymerase (Invitrogen™, Carlsbad, CA, USA), in a 25 μL reaction volume. The PCR reaction consisted of 0.25 μL Taq Polymerase, 2.5 μL of 10X buffer (Invitrogen™, Carlsbad, CA, USA), 0.16 μM of each primer, 0.16 mM of dNTP mix, 2 mM MgCl_2_ and 25 ng of template DNA. PCRs were carried out with an initial denaturing step at 95°C for 2 min followed by 30 cycles of denaturing at 95°C for 45 sec, annealing of primers at 59°C (62°C for *Forl*_155.3) for 45 sec and primer extension at 72°C for 45 sec. The PCR was completed by a final extension at 72°C for 10 min. PCR products were purified using a QIAquick PCR Purification Kit (Qiagen) and then sequenced by Sanger platform.

EF1a sequences obtained from 22 out 36 of *Foph* related isolates, were submitted to the GenBank with accession numbers (MT738937-MT738958). A total of 29 EF1a sequences of *Foph* related strains were aligned (MUSCLE method) using MEGA version 7 (Kumar et al., 2016). The corresponding EF1a sequence from the selected *F. oxysporum* ff. spp. mentioned above, were also included for comparison. Phylogenetic analysis was performed using the software BEAST (Bayesian Evolutionary Analysis Sampling Trees) v 2.6.1 (Bouckaert et al., 2019), with default settings. The resulting phylogenetic trees were visualized using the Interactive Tree of Life (iTOL) v4 (Letunic and Bork, 2019). The EF1a from *F. fujikuroi* was used as an outgroup. To corroborate the presence of the new effectors in the *Foph* related strains, specific primers for the new candidates were designed and used for PCR amplification (Supplementary Table S3), using the same conditions as mentioned above. DNA from Colombian strains of *Fol, Foc* R1 and TR4, were provided by Dr Mauricio Soto (AGROSAVIA), and used as a control for amplification.

## Results

### *Foph* genome sequencing and assembly

The genome sequence of the highly virulent strain of *Foph* (MAP5) in cape gooseberry was assembled from 250 bp paired end reads Illumina MiSeq into 1856 contigs with a total size of 44.9 Mb. This genome assembly is smaller, compared to the reference genomes of different *ff. spp* of *F. oxysporum* (ranging from 47 to 61 Mb approximately), specific for *Solanaceous* and *Alliaceous* hosts, other Illumina genome assemblies available for strains grouped in the f. sp. named *Fophy*, that infect other *Physalis* host species (i.e. husk tomato or *P. philadelphica*), and two strains that infect tobacco (Fonic_003) and eggplant (Fomel_001) respectively. (Ma et al., 2010; van Dam et al., 2016, 2017; Armitage et al., 2018). Despite its fragmentation, the predicted gene content of this genome assembly of *Foph* (15019 transcripts), is similar to the illumina genome assemblies available in the GenBank (Table 1, Supplementary Table S1).

**Table 1.** *Foph* genome assembly statistics, compared to other Illumina genome sequences of ff. spp. of *F. oxysporum* Solanaceous infecting strains and two nearly complete genome assemblies of *Fol* and *FoC.*

| Strain | Seq. Platform | No. of Contigs | Maximum length (kb) | N50 (kb) | GC (%) | Assembly length (Mb) | Transcripts |
| --- | --- | --- | --- | --- | --- | --- | --- |
| *Foph* | Illumina | 1856 | 453 | 70 | 48.5 | 44.9 | 14897 |
| **MAP5** |  |  |  |  |  |  |  |
| *Fophy* | Illumina | 488 | 2037 | 1167 | 47.7 | 47.2 | 23095 |
| **KOD886** |  |  |  |  |  |  |  |
| *Fophy* | Illumina | 1275 | 1667 | 547 | 47.6 | 50.4 | 24279 |
| **KOD887** |  |  |  |  |  |  |  |
| *Fomel* | Illumina | 1725 | 2348 | 227 | 47.5 | 52.3 | 16492 |
| **001** |  |  |  |  |  |  |  |
| *Fonic* | Illumina | 638 | 2572 | 1159 | 47.6 | 49.9 | 15480 |
| **003** |  |  |  |  |  |  |  |
| *Fol_4287* | PacBio | 88 | 6854 | 458 | 48.3 | 61.4 | 27347 |
| *FoC_Fus* | PacBio | 34 | 6434 | 414 | 47.7 | 53.4 | 19342 |

### Comparative genomics of *Foph*

A total of 14897 transcripts were predicted from the genome assembly of *Foph*, from which 14140 have an orthologous counterpart in the genomes of *F. oxysporum* compared in this study (Table 1). Using the anvi’o pipeline for pangenome analysis, a set of the single copy orthologous genes (SCG) present in the 22 *F. oxysporum* genomes were extracted to reconstruct their phylogenetic relationship. We used this phylogenetic reconstruction to test whether *Foph* could be related to *Fophy* (i.e. *Physalis* infecting strains) or might be grouped in a lineage of strains that infect *Solanaceous* hosts. The resulted phylogenomic tree showed that *Foph* shared the same clade with *Fonic* and *Fol_R3* and is closely related to *Forl* and *Fo47* (both strains associated to the tomato crop). Nevertheless, no closer relationship was found between *Physalis* infecting ff. spp. *(Foph* and *Fophy*), indicating their polyphyletic origin and different host specificity (Figure 1a). We also performed a comparative analysis using the SCG shared between *Foph* and the remaining 21 genomes of *F. oxysporum* ff. spp. As expected, this analysis showed that the majority of *Foph* SCG (~14K), are syntenic with the core chromosomes of *Fol* (used here as the reference genome sequence of *F. oxysporum* species). These syntenic SCG might correspond to the core genome of *Foph*, while the remaining ~0.5K of *Foph* SCG, correspond to transcripts that are not present in any cluster and could be part of the LS genomic regions (Figure 1b).

**Figure 1.**
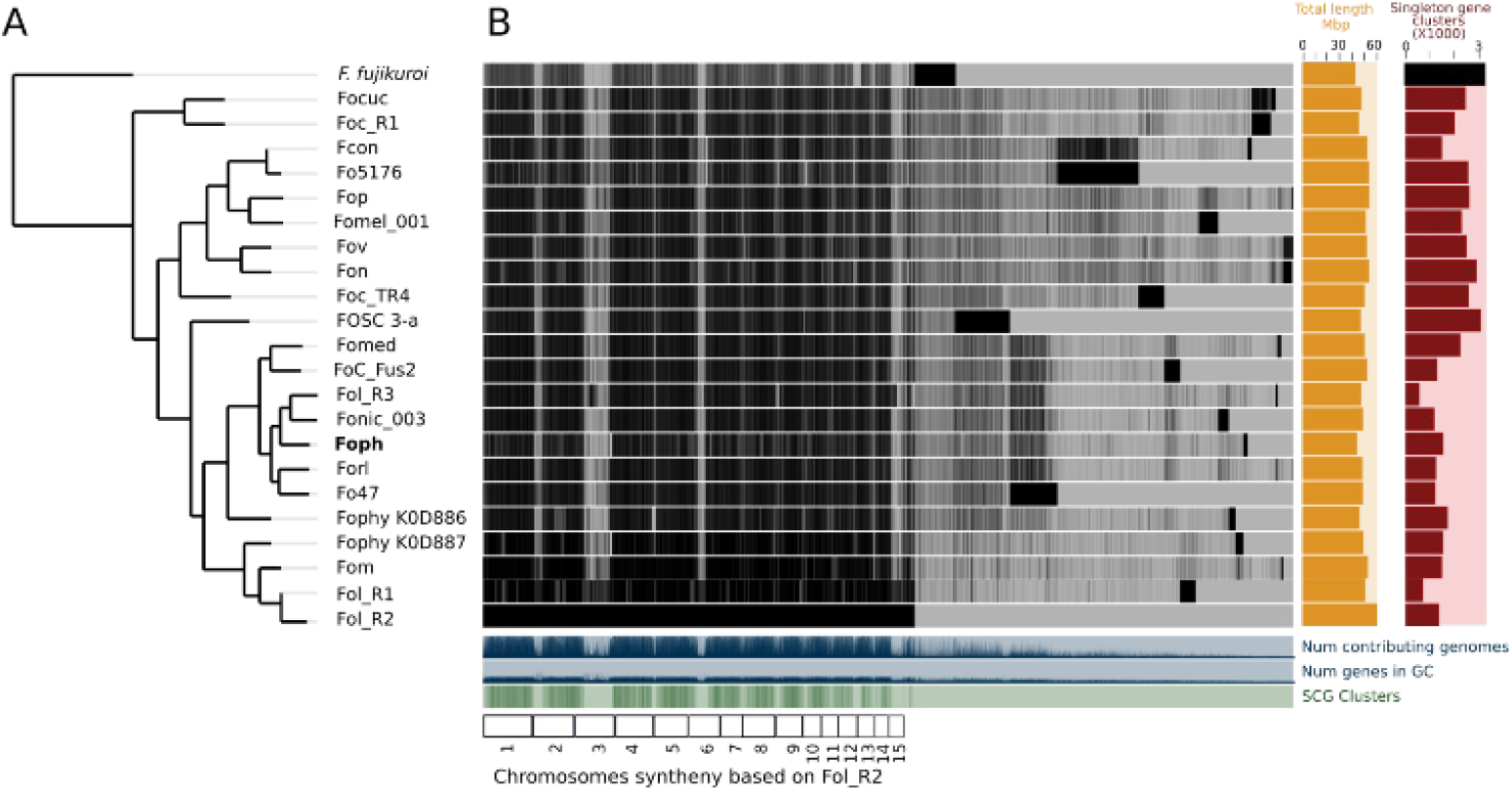
Comparative genomics between *Foph* and 21 *F. oxysporum* ff. spp. **A.** The phylogenomic tree was inferred with the single copy orthologous genes of the 22 genomes of *F. oxysporum* used in this analysis. The amino acid sequence of the translated genes was concatenated, and the final alignment consists of a total of 4184361 amino acid positions. The phylogenetic tree was constructed using FastTree 2.1. *F. fujikoroi* (IMI58289), was used as outgroup. **B**. Pan-genomic analysis of *F. oxysporum*, showing the core genome the species complex and single copy orthologous genes, possibly forming the LS genome for each forma specialis. The *F. oxysporum* pan-genome was generated using the anvi’o pangenomic workflow.

### The homologous effectors are confirmed in the genome sequence of *Foph*

In our previous study, eight homologous effectors were identified in *Foph* by i*n planta* RNAseq mapping analysis with the LS regions of *Fol* (Simbaqueba et al., 2018). Here, we performed a combination Blastp and Blastx searches of the known SIX effectors and Ave1 effectors in the genome assemblies of the 24 *F. oxysporum* ff. spp. (including *Foph*), compared in this study (Table 2). This result showed the widespread presence of SIX homologues in different ff. spp. of *F. oxysporum* and specifically, confirming the presence of highly identical (87 to 100 %) of *Fol* homologous effectors in the genome of *Foph.* Interestingly, in this search we also identified a highly identical putative homologous transcript of the *Fol* SIX13 effector present in *Foph.* This prediction was manually confirmed as the corresponding transcript of SIX13, was fragmented into two contigs (ctg_1292 and ctg_1535) in the *Foph* genome (Table 3).

**Table 2.**
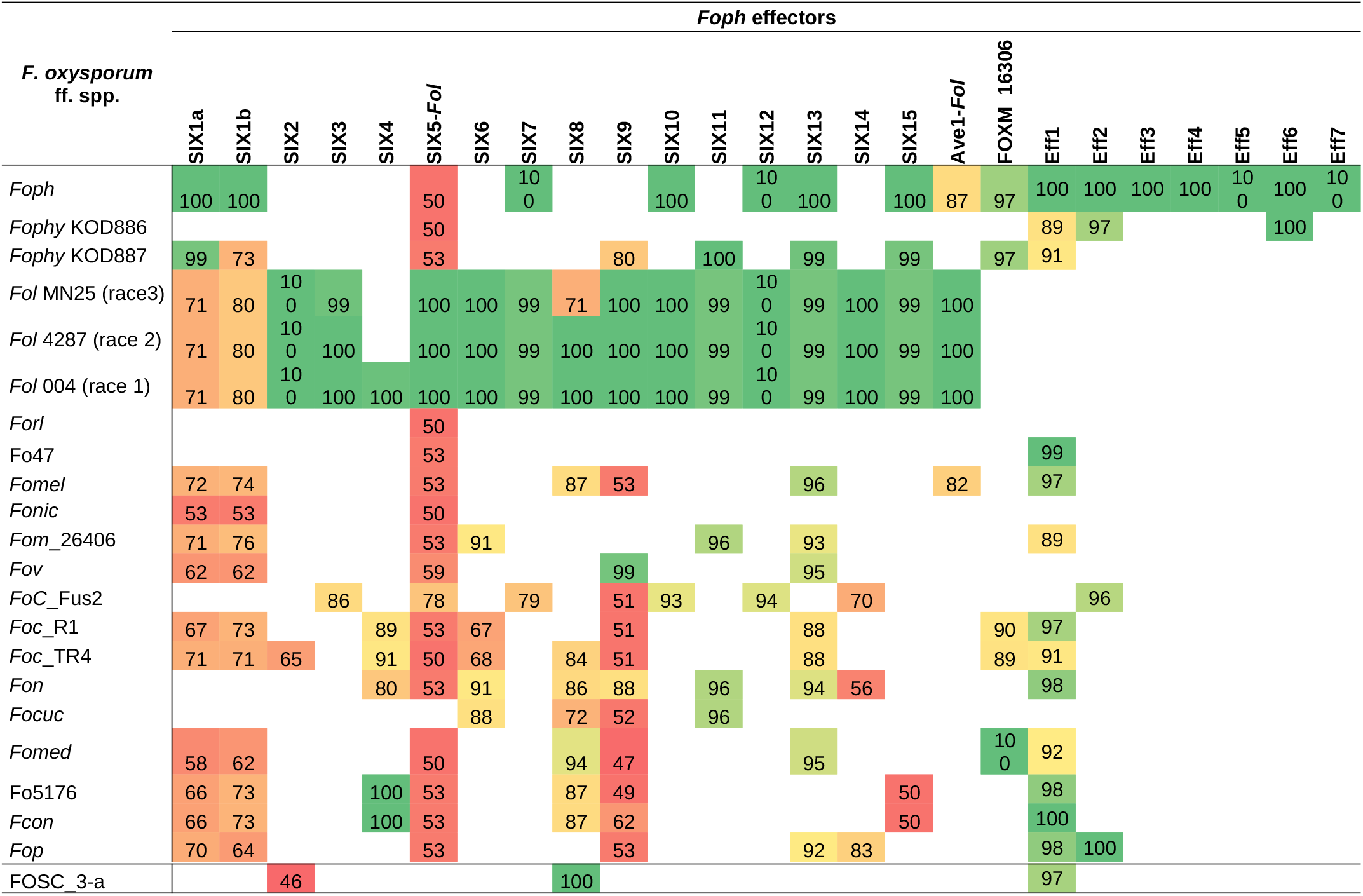
tBlastn identities of *Foph* effectors compared against the *F. oxysporum* species complex WGS databases

**Table 3.**
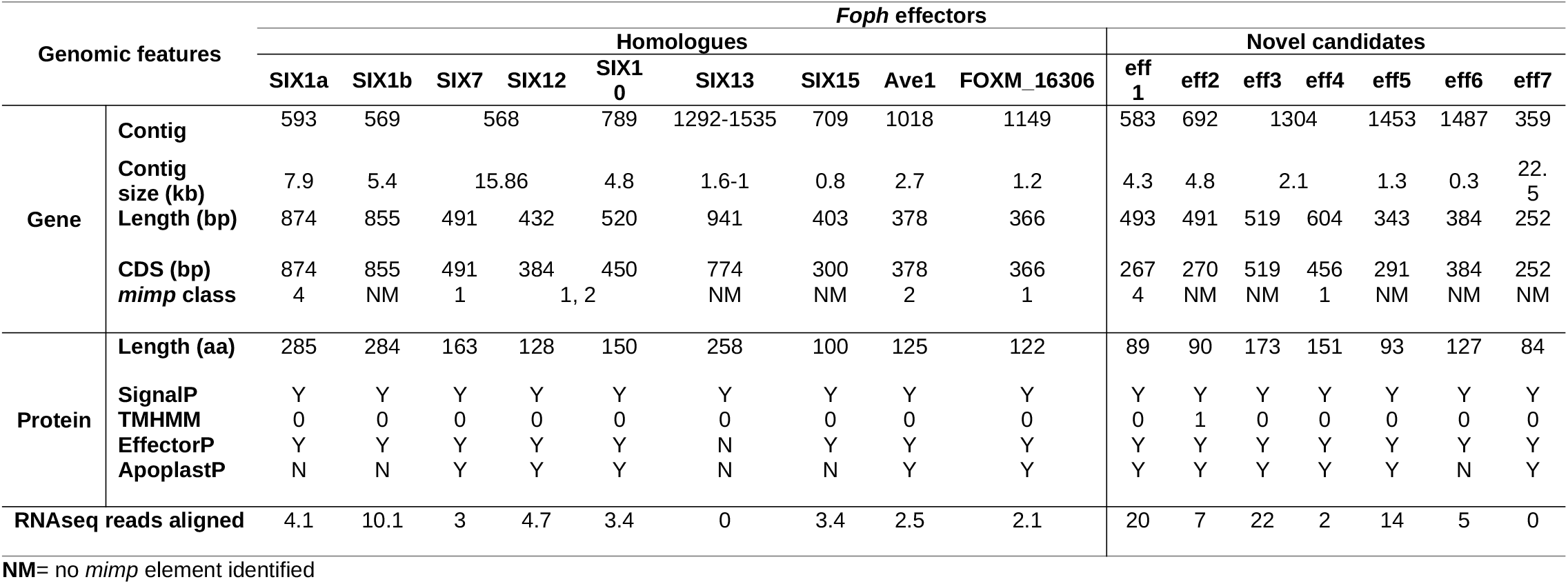
Genomic analysis of the effectors identified in the *Foph* genome

These results also confirmed that the *Fol* effector gene cluster formed by the SIX7, 12 and 10 (Ma et al., 2010; Schmidt et al., 2013) and partially identified in *Foph* by *in planta* transcriptomics (Simbaqueba et al., 2018), is entirely conserved in the genome of *Foph.* SIX7 and SIX12 homologues are both present in the same contig (ctg_568) while SIX10 is located in another contig (ctg_789) of the *Foph* genome assembly (Table 3). Thus, we manually inspected the sequences of these contigs and found that both contigs are overlapped by a sequence segment of 22 bp at the proximal 5’ end of the ctg_586 with the distal 3’ end of the ctg_789. This overlapped segment of both contigs correspond to a *mimp* class 2 sequence in intergenic region between SIX10 and SIX12. The *Foph* effector gene cluster is 4.7 kb in length and is similar to that formed by the same homologous effectors in *Fol* (5.2kb), including the intergenic regions with the approximate same length as *Fol* (i.e. 1.8 kb between SIX7 and SIX12 and 1.4 kb between SIX12 and SIX10, respectively), and three *mimp* elements that flank the effector gene cluster reported by Schmidt et al, (2013) in *Fol* (Figure 2).

**Figure 2.**
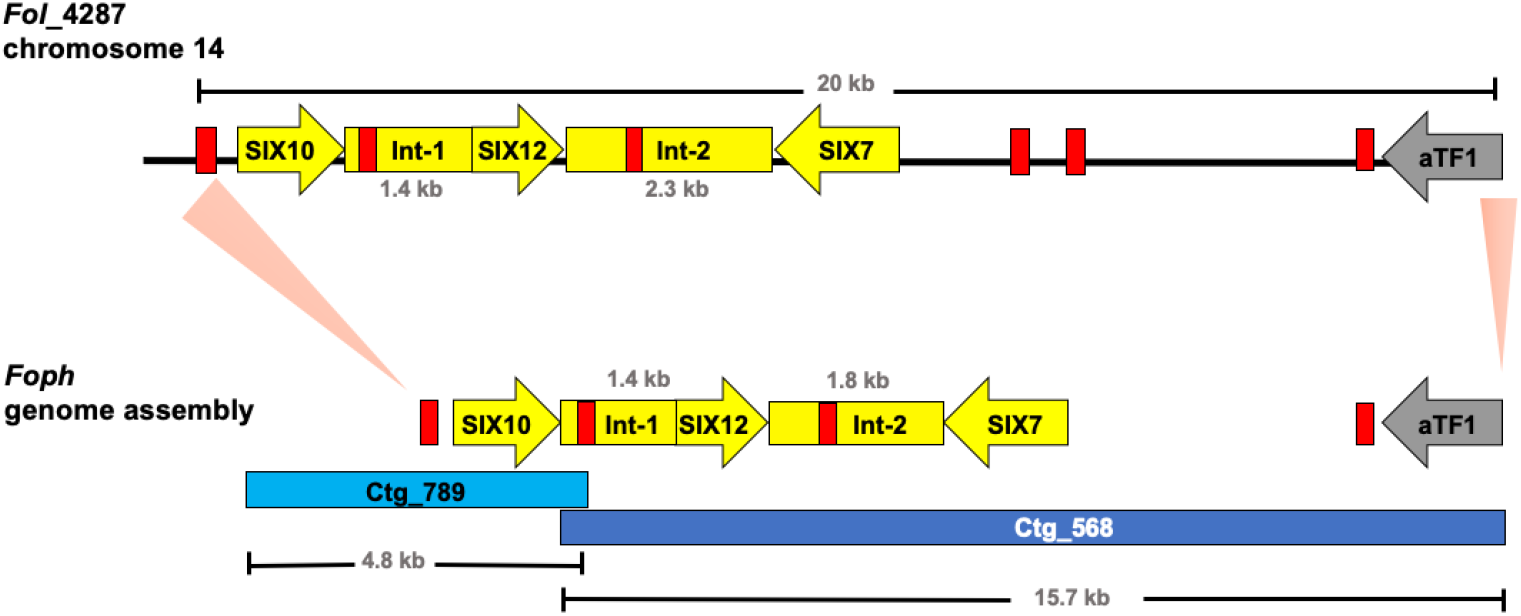
Graphical representation of a 20 kb segment of the chromosome 14 in *Fol*, containing the cluster of effectors SIX10, 12 and 7, and the aTF1 gene (FOXM_17458) (upper part). The chromosomal segment is conserved in the *Foph* genomic region (shown by orange pale triangles), corresponding to the overlapped Contigs 568 and 789 (Bottom part), suggesting a highly possible horizontal transfer of a chromosomal segment of 20kb between both ff. spp. Int-1= conserved intergenic region between SIX10 and SIX12. Int-2= conserved intergenic region between SIX12 and SIX7. Red blocks represent *Mimp* transposable elements flanking the cluster of effector genes shared between *Fol* and *Foph.*

Further inspection of the ctg_568, also confirmed the presence of another highly conserved homologous gene (FOXG_17458) between *Foph* and *Fol*, including the corresponding *mimp* class 1 element in the 5’ UTR (Figure 2). The transcript FOXG_17458 in *Fol*, encode one transcription factor of the family *a*TF1 - FTF1 (van der Does et al., 2016), and is located 9 kb away from the ORF of the SIX7, while its counterpart homologous sequence presented in *Foph* is located 7kb away from the SIX7 ORF. This finding suggests a highly probable horizontal transfer of a chromosomal segment of at least 20 kb in length between *Fol* and *Foph* (Figure 2). In *Fol*, SIX15 is a non-annotated transcript and is located 55 kb away from the *a*TF1. This chromosome region includes four annotated transcripts: FOXG_17459, FOXG_17460, FOXG_17461 and FOXG_17462. Thus, we performed a Blastn search using this sequence of 55kb from *Fol* as a query and compared with the *Foph* genome assembly, in order to test whether an extended sequence of the chromosome 14 of *Fol* is conserved in *Foph.* However, no additional chromosomal segment shared between *Fol* and *Foph* was identified by comparing both genomic sequences.

### Novel candidates for effector genes in *Foph*

We identified novel effector genes in the *Foph* genome, by combining the sets of proteins from the secretome and effectorome respectively. We predicted a total of 1495 secreted proteins, forming the *Foph* secretome, from which 276 were determined to be effectors, named herein as “*Foph* effectorome”. Seven transcripts of the *Foph* effectorome were identified as novel effectors, due to the lack (*Foph_eff2, Foph_eff3, Foph_eff4* and *Foph_eff7*) or low similarity (*Foph_eff5* and *Foph_eff6*) to any protein reported in the public databases (Table 2). Additionally, *mimp* elements were identified 624bp and 430bp upstream from the transcripts *Foph_eff2* and *Foph_eff5*, respectively (Table 3)

The candidate effector *Foph_eff1* showed significant tBLASTn hits with different *F. oxysporum* ff. spp., including the non-pathogenic Fo47. Therefore, this transcript could be excluded as a novel effector gene. The unique candidates *Foph_eff3* and *Foph_eff4* are clustered in the contig_692 at 700 bp of distance approximately between them. Furthermore, we predicted a transmembrane domain for protein encoded by *Foph_eff3* (Table 3), suggesting a cellular localization and with a possible different function from a secreted protein. Additionally, we performed an RNAseq mapping against the ORF of the novel candidate effectors and found that six out of the seven candidates are expressed during cape gooseberry infection at 4 dpi. In this analysis we also included the homologues of SIX effectors and homologues in *Foph* of the EF1a, Beta tubulin chain (*β-tubulin*) and *Fusarium* extracellular matrix 1 (*FEM1*), as housekeeping genes for expression controls. We found that *Foph_eff1*, *eff4, eff6* and *eff7*, showed higher expression compared to the rest of the transcripts analysed (Figure 3). Interestingly, *eff2, eff4*, and *eff6*, showed higher expression, compared to all three controls. These results support the evidence of these novel candidates as putative effectors in *Foph*, that could be involved in pathogenicity.

**Figure 3.**
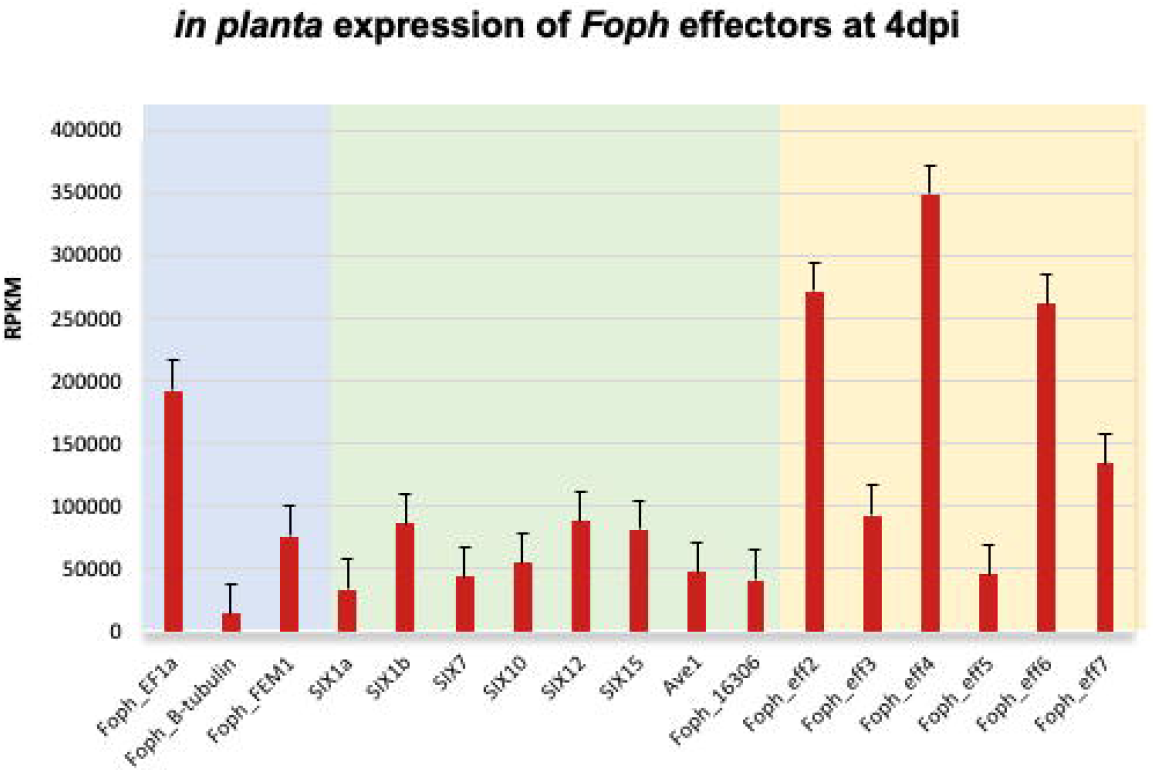
Expression analysis of the effectors identified in the genome sequence of *Foph_MAP5*, using the RNAseq data form cape gooseberry susceptible plants inoculated with *Foph* at 4 dpi, reported in our previous *in planta* transcriptomic analysis. Pale blue panel indicate the genes translation elongation factor alpha (EF1a), tubulin B-chain (B-tubulin) and *Fusarium* Extracellular Matrix 1 (FEM1), to use as constitutive expressed control genes of *Foph* during host infection. Pale green indicates the expression of the homologous effectors identified in *Foph.* Pale yellow indicates the expression of the newly identified effectors in *Foph*. Six out of seven new effector candidates are expressed during cape gooseberry infection with a higher expression of *eff2, eff4* and *eff6*, compared to the rest of the effectors analysed. RPKM= reads per kilobase per million of mapped reads. Scale bars indicate standard error

### Novel effectors are present in *F. oxysporum* isolates associated to the cape gooseberry crop

In order to test if the candidate effectors genes could be used as potential molecular makers for *Foph* identification in diagnostic strategies, we performed a preliminary screening of the novel candidate effectors by PCR amplification in a panel of 36 *F. oxysporum* isolates (including *Foph-* MAP5), obtained from cape gooseberry crops. The isolates have been classified, based on their ability to cause wilting symptoms (32) and non-pathogenic (4), on a susceptible cape gooseberry genotype (Supplementary Table S2). The screening also included DNA isolated from *Fol*, *Foc*R1 and *Foc*TR4 strains, as control for amplification. We found amplification for all candidates in the majority of *F. oxysporum* isolates associated to cape gooseberry, including pathogenic and non-pathogenic (Supplementary Table S2 and Figure S1). Therefore, we did not identify specificity of the candidate effectors for the putative *Foph* pathogenic isolates. However, we did not identify the presence of the novel effectors *Foph_eff3*, *eff4*, *eff5, eff6* and *eff7* in the control strains *Fol, FocR1 and FocTR4.* This result suggests that these five novel candidates could be specific for *F. oxysporum* strains associated to the cape gooseberry crop. We also conducted a molecular characterization using the *EF1a* sequence of 28 out of 36 *F. oxysporum*, in order to test whether these isolates associated to cape gooseberry host, might be originated from a single linage. However, the phylogenetic tree showed that these 28 isolates are grouped together in two different lineages, compared to the ff. spp. of *F. oxysporum*, suggesting the polyphyletic origin of *Foph* related strains (Figure 4).

**Figure 4.**
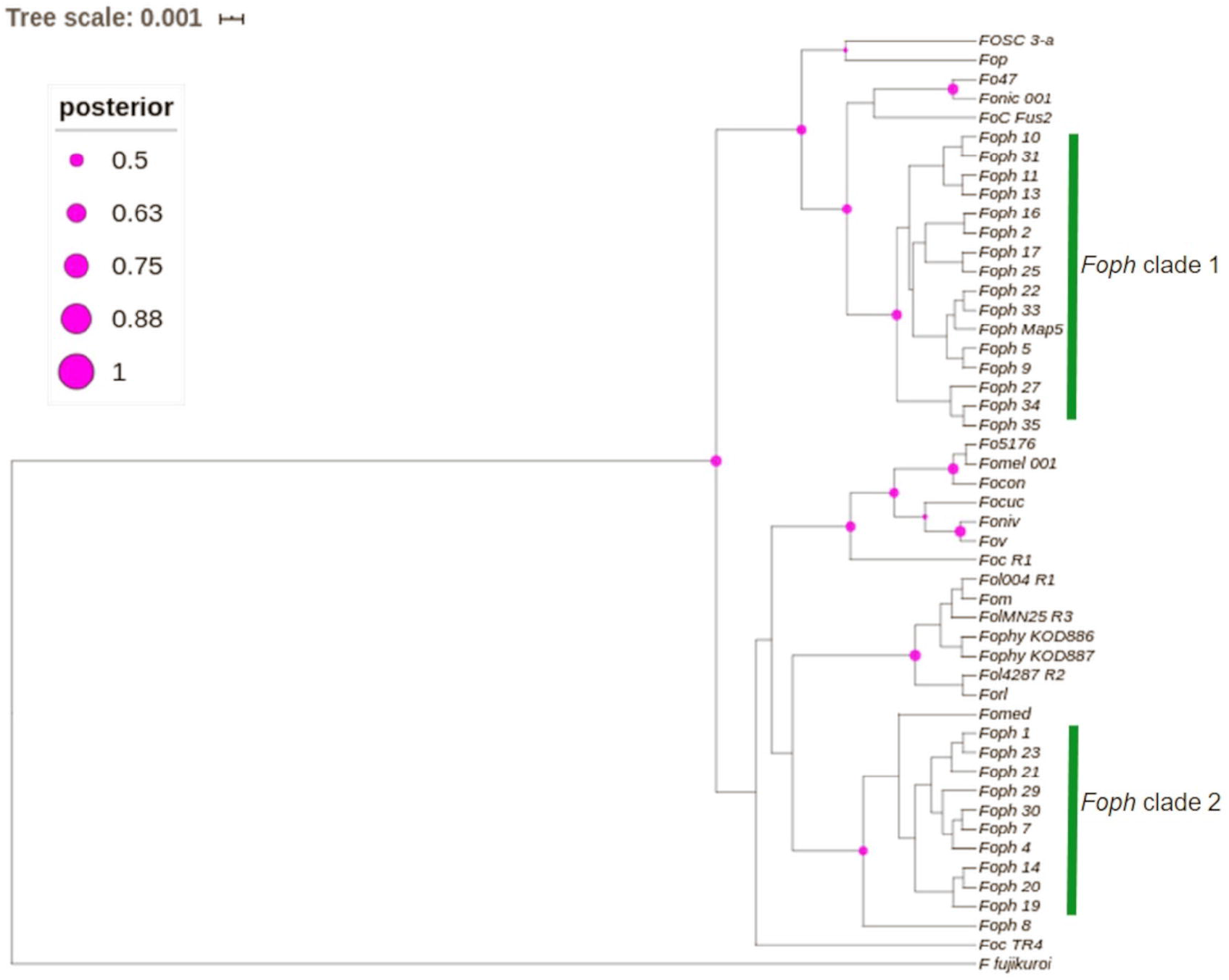
Phylogenetic tree of a partial sequence of the EF1a gene from the genome sequences of 24 ff. spp. of *F. oxysporum* (Table S1) and from 26 *F. oxysporum* isolates obtained from cape gooseberry crops (Table S2). The phylogenetic analysis was conducted using BEAST. Node shapes indicate the bootstrapping support, indicated as Bayesian posterior probabilities. The scale bar indicates time in millions of years.

## DISCUSSION

### *Foph* genome and phylogenetic relationship with other ff. spp

In Colombia, the cape gooseberry crop is severely affected by pathogenic strains of *Foph*, with losses of nearly 90%. In this pathosystem, SNPs associated to resistant cape gooseberry genotypes, *Foph* pathogenic strains and homologous effectors have been identified (Osorio-Guarin et al., 2016; Simbaqueba et al., 2018). However, there is a need to implement genomic approaches to corroborate these findings and to identify new sources associated to the interaction between *Foph* and cape gooseberry. These approaches could be used in the development of disease management strategies and plant breeding programs in the cape gooseberry crop. Here we sequenced and assembled the genome of the highly virulent strain *Foph*-MAP5, aiming to identify novel candidates for effector genes that could be characterized in further studies and implemented in diagnostic strategies. Comparative and functional genomics of F. oxysporum that infect cucurbit species, suggested that their host range could be determined by the close phylogenetic relationship associated to their homologue effector gene content (van Dam et al., 2016, 2017b). This hypothesis is supported by additional evidence on the formae speciales *radicis-cucumerinum (Forc*) and *melonis* (*Fom*), showing that a syntenic LS chromosome region is highly related to the expansion formae speciales range (van Dam et al., 2017b; Li et al., 2020b). Recent genome analysis on the chromosome-scale assembly of the brassicas infecting f. sp. Fo5176, showed a similar pattern of phylogenetic relationship possibly associated to the expansion of their host range (Fokkens et al., 2020). We performed comparative genomics using the *Foph* genome assembly in order to test whether a set of available genomes of Solanaceous-infecting formae speciales including *Foph*, could show a similar phylogenetic related pattern. Nevertheless, our analysis showed that the tested strains have a different ancestry (Figure1), despite the close relationship of *Foph* with tomato infecting ff. spp. *Fol*-R3, Fo47, Forl and tobacco Fonic_003, and our previous evidence on horizontal gene transfer of effectors between *Fol* and *Foph.* Resequencing of the genomes including *Foph, Fophy*, *Fonic* and *Fomel*, using long reads, will help to gain a deeper understanding of the phylogenetic relationship among Solanaceous-infecting ff. spp.

### Confirmation of homologues and identification of new ones

Homologues of *Fol SIX* genes have been identified in other ff. spp. of *F. oxysporum* and other *Fusarium* species (Thatcher et al., 2012; Meldrum et al., 2012; Rocha et al., 2016; Schmidt et al., 2016; Li et al., 2016; Taylor et al., 2016, 2019; van Dam et al., 2016, 2017a; Williams et al., 2016; Simbaqueba et al., 2018; Armitage et al., 2018). The presence of the SIX homologues might be a consequence of horizontal transfer of genes or segments of pathogenicity chromosomes between different strains of *F. oxysporum* and/or fungal phytopathogenic species. In our previous study, we identified homologues of the SIX, Ave1 and FOXM_16306 effectors, analysing an in *planta* RNAseq of *Foph.* Despite the fragmentation of this genome assembly (i.e. no scaffolds generated), we corroborated the presence of complete sequences of the homologous effectors SIX, Ave1 and FOXM_16306, contained in different contigs that could correspond to the LS regions of the *Foph* genome (Tables 2 and 3).

We also found a homologue transcript of the *Fol* SIX13 in the genome of *Foph*, fragmented into two contigs. This homologue was not expressed at 4 dpi and therefore, it was not identified in our previous transcriptomics study. SIX13 homologues are present in legume, cucurbits, musaceous and solanaceous infecting ff. spp. of *F. oxysporum* (Ciszlowski et al., 2016; van Dam et al., 2016; Williams et al., 2016). The later mentioned ff. spp., are highly identical at the protein level (96% in *Fomel* and 99% in *Foph* and *Fophy*, respectively) (Table 2). In cucurbits infecting ff. spp. of *F. oxysporum*, a suit of effectors was found to be associated with host specificity (van Dam et al., 2016). Thus, the highly identical SIX13 homologues in the Solanaceous-infecting ff. spp., could be related to their specificity for these group of host species. Moreover, the majority of the SIX genes in *Fol* are located on the chromosome 14 (i.e. pathogenicity chromosome), except for SIX13, which is found in the LS chromosome 6 (Schmidt et al., 2013). Similarly, SIX13 corresponding homologues of *Fomed* and *Foph* are located on LS regions (Williams et al., 2016; Table 2). In *Foc*, SIX13 homologues, have been associated to the differentiation of TR4 and R4 and are currently used in molecular based diagnostic of TR4 in banana crops (Cahrvalis et al., 2019). Together, this evidence suggests that SIX13 could play a role in pathogenicity or host specificity. Future functional analysis of on *Foph*-SIX13 is necessary to confirm this hypothesis.

Furthermore, we performed a manual inspection of the contigs 568 and 789 of the *Foph* genome and confirmed the presence of a highly conserved chromosomal segment of 20kb of *Fol* that includes a cluster of physically linked effector genes (SIX7, SIX10, SIX12 and extended transcription factor αTF1). This shared region also included their corresponding flanking *mimp* elements (Figure 2, Schmidt et al., 2013; Simbaqueba et al., 2018). This finding suggests a highly probable horizontal acquisition of an entire genomic segment of 20kb from an ancestor of *Fol* or *Foph.* Miniature impala (*mimp*) transposable elements (TEs), have been identified in the genome sequences of different phytopathogenic fungi of the *Fusarium* genus (Schmidt et al., 2013; van Dam and Rep, 2017). In *F. oxysporum*, *mimp* elements have been associated to the gain or loss of effector genes, presumably acting as an evolutionary mechanism of emergence of new phytopathogenic strains (van Dam et al., 2017b). The presence highly identical *mimp* elements, flanking the homologous effector gene cluster in both *Fol* and *Foph* (Figure 2), suggests that these TEs could play a role in the lateral transference of this homologue genomic region between *Foph* and *Fol*.

Functional analysis of SIX effectors in *Fol*, showed that mutant strains with a large deletion (0.9 Mb) of chromosome 14, including the candidate effector genes SIX6, SIX9 and SIX11 did not show any loss of virulence compared to wild type *Fol* on tomato plants (Vlaardingerbroek et al., 2016). Recent evidence revealed by another set of *Fol* mutant strains with chromosomal deletions that include the SIX10, SIX12 and SIX7 gene cluster, showed no loss of virulence on tomato plants (Li et al., 2020a). These findings indicate that the genes located in these chromosomal segments (including the SIX genes with homologues in *Foph*), could be dispensable for pathogenicity, while the remaining segments could be sufficient for tomato infection (Vlaardingerbroek et al., 2016; Ling et al., 2020a). Although neither of the SIX7, SIX10 and SIX12 effector genes have a role in *Fol* virulence, the presence of the highly identical homologues between *Fol* and *Foph*, suggests that this segment could be undergoing adaptation to another environment (i.e. a different host plant). Therefore, it might be possible that SIX7, SIX10 and SIX12 have a role in *Foph* pathogenicity. Future investigation about the function of this conserved genomic region between these two Solanaceous-infecting ff. spp., is required. Crossed pathogenicity assays inoculating tomato and cape gooseberry with *Fol* and *Foph* and knock out of the gene cluster in *Foph* could be performed to support these hypotheses.

In this study, we confirmed that homologues of Ave1 have been only identified in the solanaceous infecting ff. spp. *Fol, Foph, Fomel*001 (Table 2) and in the f. sp. *gladioli* of *F. oxysporum* (Simbaqueba et al., 2018). Ave1 could also be present in putative conditional dispensable segments on the *Foph* genome (Table 2, Figure 1). The presence of less conserved homologues of *Fol* including SIX1 and Ave1, which are also located on *Fol* chromosome 14 (Schmidt et.al., 2013), suggests that these effectors may have a different ancestry, via acquisition of different segments of the pathogenicity chromosome at different times in the evolution of *Fol* or *Foph*.

In the tomato pathogen *Verticillium dahliae, Ave1* is involved in pathogenicity, while there is no evidence that its homologue present in *Fol* has a role in virulence (de Jonge et al., 2012; Schmidt et al., 2013). Furthermore, *Fol-*Ave1 is not expressed during tomato infection (Catanzariti, personal communication). Conversely, we found that *Foph-*Ave1 was expressed during cape gooseberry infection (Figure 3). This finding suggests that *Ave1* might have a role in *Foph* pathogenicity. Therefore, functional analyses are required by generating gene knockout strains in *Foph.* In both *V. dahliae* and *Fol, Ave1* could act as avirulence factors since they are recognised by the tomato receptor Ve1 (de Jonge et al., 2012). The *Ave1* homologue of *Foph* is highly similar at the protein level to its counterparts in *F. oxysporum* (*Fol*, *Fomel* and *Fogla*), and less similar to *V. dahliae* Ave1 (Table 2, Simbaqueba et al., 2018). The presence of Ave1 in *Foph*, suggests that the avirulence function of *Fol* Ave1 might be conserved. This hypothesis needs further investigation e.g. by testing for recognition of *Foph* Ave1 by tomato *Ve1* or a homologue in cape gooseberry.

Novel candidate effectors in *F. oxysporum* have been reported for other ff. spp., including *Fom*, *Foc_*Fus2, *Fonar* and legume infecting strains (Schmidt et al., 2015; Taylor et al., 2016, 2019; Williams et al., 2016; Armitage et al., 2018; van Dam et al., 2018), based on the analysis of their genome sequences to identify transcripts that encode for small proteins with a secretion signal peptide and the proximity of mimp to the start codon. Here, we used the predicted transcripts from the genome assembly of *Foph* to identify novel effectors, based on the effectorome, secretome repertoires and the absence or low similarity to any predicted or non-predicted protein sequences compared to genomes available in the public databases (Tables 2 and 3). Three highly expressed novel effectors during infection (*Foph*_*eff2*, *eff4* and *eff7*), are unique *Foph* candidate effectors, while the other highly expressed candidate *Foph_eff6*, have identical homologous proteins in the genomes of *Fomel* and *FoC*_Fus2 (Table 2). Furthermore, the homologous counterpart identified in *FoC_*Fus2 is located in a lineage specific region (Armitage et al., 2018). These findings suggest that *Foph_eff6* and its homologues, may have a putative role in pathogenicity and represent a subject for future functional analysis.

### Presence of effectors in the *Foph* strains and compared to other ff. spp

*Foph* pathogenic strains are responsible for the wilting disease that affect cape gooseberry crops in Colombia. Thus, appropriate disease management strategies are needed to be implemented (Barrero et al., 2012). However, the development of those strategies has been largely limited due to the lack of knowledge of the wilting disease caused by *Foph*, and accurate identification of pathogenic strains. Detection methods based on the use of effector genes as molecular markers are highly desirable for precise identification of pathogenic strains in disease management programs of soilborne pathogens due to their limited sequence diversity between members of the same *f. sp.* (Rocha et al., 2016; Gordon TR, 2017), thus providing a solid and sensitive identification of pathogenic strains of soilborne pathogens including *F. oxysporum* (van Dam et al., 2017a; Cahrvalis et al., 2019; Taylor et al., 2019).

Comparative genomics have been performed to design molecular markers based on candidate effector genes and successfully tested for the identification of cucurbit and *Narcissus* Infecting ff. spp of *F. oxysporum.* (van Dam et al., 2016; Taylor et al., 2019). In this study, we used the highly conserved novel candidate effectors found by comparative genomics in *Foph*, to explore their usefulness as potential molecular markers specific for pathogenic strains. The presence of homologous effectors suggests a functional redundancy between different ff. spp. (Taylor et al., 2019). Here, we identified that the candidate novel effector *Foph_eff1* has homologues in other ff. spp. (Table 2). We also identified the presence of *eff1* in all tested strains, including *Fol* and *Foc.* Thus, the role of *eff1* in pathogenicity may be dispensable due to its presence in different *F. oxysporum* strains and could be discarded for diagnostic purposes. The remaining novel effectors showed a clear pattern of amplification in *F. oxysporum* strains associated to the cape gooseberry crop, compared to the highly pathogenic *Fol* and *Foc* in tomato and banana respectively (Supplementary Figure S1 and Table S2). However, we did not find an amplification pattern associated to the pathogenic strains for any of the effectors tested. A similar inconsistent pattern of presence/absence between pathogenic and non-pathogenic cucurbit infecting strains of *F. oxysporum* was observed for some of the effectors-based markers developed by van Dam et al, (2017a). These results might be supported by the fact that effectors show limited sequence diversity between strains of the same f. sp. (van Dam et al., 2017; Taylor et al., 2019). An alternative explanation could be related with the limited number of effectors-based markers identified in this fragmented genome assembly of *Foph.* New markers associated to *Foph* pathogenicity will be predicted in future studies, enlarging effectorome repertoire from the resequencing of the *Foph* genome using long reads sequencing technologies as performed for *Forc* and *FoC_Fus2* (van Dam et al., 2017b; Armitage et al., 2018).

## Data availability statement

The genome assembly of *Foph* is available on NCBI under the BioProject accession number PRJNA640423. GenBank accession numbers: MT738929 – *Foph* SIX13, MT38930-MT38936-*Foph* eff1 to eff7, MT738937 - MT738958 EF1a sequences of *F. oxysporum* strains associated to cape gooseberry crops. Access to these sequences must be requested to the Ministry of Environment and Development of Colombia.

*Foph* strains used in this work were collected under the framework collection permit No.1466 from 2014 of AGROSAVIA and registered in the National Collections Registry (RNC129) of Colombia

## Author Contributions

JS planned and carried out the *Foph* genome analysis, planned the experiments, analysed the data, created figures, and drafted, wrote and edited the manuscript. ER and DB carried out the experiments with *Foph* isolates. CG obtained funding, planned experiments, contributed and edited the manuscript. AC obtained funding, planned and carried out the *Foph* genome sequencing, analysis and all bioinformatics, created figures, drafted and edited the manuscript.

## Acknowledgments

J.S. was supported by a Postdoctoral Fellowship from the Ministry of Science, Technology and Innovation (MINCIENCIAS), Colombia. We thank to Johan David Barbosa for his contribution to the results obtained in the molecular and pathogenic characterization of *Foph* strains, reflected in the Supplementary Table 2. We thank to Dr Mauricio Soto (AGROSAVIA) for provided DNA from Colombian strains of *Fol, Foc R1* and *TR4*, used as PCR amplification controls. We thank grateful to Ministry of Agriculture and Rural Development of Colombia for the financial support.

