## Supplementary Table 1 for "Novel effector genes revealed by the genomic analysis of the phytopathogenic fungus *Fusarium oxysporum* f. sp. *physali* (*Foph*) that infects cape gooseberry plants"

**Table S1.** Description of the *F. oxysporum* ff. spp. and *F. fujikuroi* used in this study for comparative genomics

| **GeneBank accession** | ***forma specialis*** | **Abbreviation**  **(f. sp.)** | **Strain/isolate** | **Host** |
| --- | --- | --- | --- | --- |
| JABXJO000000000 | *physali* | *Foph* | MAP5 | cape gooseberry |
| GCA_002233995.1 | *physalis* | *Fophy* | KOD886 | husk tomato |
| GCA_002233985.1 | *physalis* | *Fophy* | KOD887 | husk tomato |
| GCA_000260495.2 | *melonis* | *Fom* | 26406 | melon |
| GCA_000260175.2 | *vasinfectum* | *Fov* | 25433 | cotton |
| GCA_000260155.3 | *radicis-lycopersici* | *Forl* | 26381 | tomato |
| GCA_002234045.1 | *nicotiane* | *Fonic*_003 | 10913 | tobacco |
| GCA_003615085.1 | *cepae* | *FoC*_Fus2 | Fus2 | onion |
| GCA_000350345.1 | *cubense* | *Foc*_R1 | race 1 | banana |
| GCA_000260195.2 | *cubense* | *Foc*_TR4 | 54006 | banana |
| GCA_001702505.1 | *niveum* | *Fon* | Fon005 | watermelon |
| GCA_001702495.1 | *cucumerinum* | *Focuc* | Foc013 | cucumber |
| GCA_001652425.1 | *medicaginis* | *Fomed* | Fom-5190a | Medicago |
| GCA_002234085.1 | *melogenae* | Fomel_001 | J-71 | eggplant |
| GCA_000222805.1 | Fo5176 | Fo5176 | Fo5176 | arabidopsis |
| GCA_000260215.2 | *conglutinans* | *Fcon* | 54008_race2 | cabbage |
| GCA_000260075.2 | *pisi* | *Fop* | HDV274 | pea |
| GCA_000271745.2 | Human FOSC 3-a | FOSC_3-a | FOSC_3-a | human (clinical) |
| GCA_000271705.2 | Fo47 | Fo47 | Fo47 | tomato |
| GCA_000259975.2 | *lycopersici* | *Fol*_R3 | MN25-race 3 | tomato |
| GCA_000149955.2 | *lycopersici* | *Fol*_R2 | 4287-race 2 | tomato |
| GCA_001702875.1 | *lycopersici* | *Fol*_R1 | Fol004-race 1 | tomato |
| GCA_900079805.1 | *Fusarium fujikuroi* | IMI 58289 | IMI 58289 | rice |
