## Supplementary Table 2 for "Novel effector genes revealed by the genomic analysis of the phytopathogenic fungus *Fusarium oxysporum* f. sp. *physali* (*Foph*) that infects cape gooseberry plants"

**Table S2.** PCR analysis of the seven *Foph* candidates of novel effectors in a panel of 36 *F. oxysporum* isolates associated to cape gooseberry crops. *Fol* and *Foc* Colombian strains used as a control

| **Department** | **Town** | **Strain** | **code** | **Pathogenicity** | **eff1** | **eff2** | **eff3** | **eff4** | **eff5** | **eff6** | **eff7** | **GenBank EF1a** |
| --- | --- | --- | --- | --- | --- | --- | --- | --- | --- | --- | --- | --- |
| **Boyacá** | Arcabuco | B52 | 1 | P | + | + | + | + | - | + | + | MT738937 |
| **Cundinamarca** | Granada | C123 | 2 | P | + | + | + | + | - | + | - | MT738945 |
| **Antioquia** | La Union | A71 | 3 | P | - | + | + | + | + | + | + | NA |
| **Boyacá** | Ciénaga | B130 | 4 | P | + | + | + | + | + | + | - | MT738954 |
| **Cundinamarca** | Granada | C18 | 5 | P | + | + | + | + | + | + | - | MT738955 |
| **Boyacá** | Arcabuco | B51 | 6 | P | + | + | + | + | + | + | + | NA |
| **Cundinamarca** | Silvania | C67 | 7 | P | + | + | + | + | + | + | + | MT738956 |
| **Boyacá** | Arcabuco | B120 | 8 | P | + | + | + | + | + | + | + | MT738957 |
| **Boyacá** | Ciénaga | B97 | 9 | P | + | + | + | + | + | + | + | MT738958 |
| **Cundinamarca** | Granada | C62 | 10 | P | + | + | + | + | - | + | + | MT738938 |
| **Antioquia** | San Pedro | A77 | 11 | P | + | + | + | + | - | + | + | MT738939 |
| **Cundinamarca** | Granada | C124 | 12 | P | + | + | + | + | + | + | + | NA |
| **Boyacá** | Arcabuco | B04 | 13 | P | + | + | + | + | + | + | + | MT738940 |
| **Cundinamarca** | San Bernardo | C100 | 14 | P | + | + | + | + | + | + | + | MT738941 |
| **Antioquia** | La Union | A133 | 15 | P | + | + | + | + | + | + | + | NA |
| **Cundinamarca** | San Bernardo | C101 | 16 | P | + | + | + | + | + | + | + | MT738942 |
| **Boyacá** | Ventaquemada | B113 | 17 | P | + | + | + | + | + | + | + | MT738943 |
| **Antioquia** | San Vicente | A135 | 18 | P | - | + | + | + | + | + | + | NA |
| **Boyacá** | Ventaquemada | B110 | 19 | P | + | + | + | + | + | + | - | MT738944 |
| **Cundinamarca** | Granada | C28 | 20 | P | + | + | + | + | + | + | + | NA |
| **Antioquia** | San Pedro | A136 | 21 | P | + | + | + | + | + | + | + | MT738946 |
| **Cundinamarca** | San Bernardo | C24 | 22 | P | + | + | + | + | + | + | + | JX465126.1 |
| **Antioquia** | El penon | A47 | 23 | NP | + | + | + | + | + | + | - | MT738947 |
| **Antioquia** | La Union | A132 | 24 | P | + | + | + | + | + | + | + | NA |
| **Antioquia** | El penon | A48 | 25 | P | + | + | + | + | + | + | + | MT738948 |
| **Cundinamarca** | Granada | C61 | 26 | P | + | + | + | + | + | + | + | NA |
| **Boyacá** | Arcabuco | B02 | 27 | P | + | + | - | - | - | - | - | JX465108.1 |
| **Cundinamarca** | Granada | C33 | 28 | NP | + | + | + | + | + | + | + | NA |
| **Cundinamarca** | Granada | C20 | 29 | P | + | + | + | + | + | + | + | JX465114.1 |
| **Boyacá** | Arcabuco | B05 | 30 | P | + | + | + | + | + | + | + | MT738950 |
| **Boyacá** | Arcabuco | B06 | 31 | P | + | + | + | + | + | + | - | MT738951 |
| **Boyacá** | Arcabuco | B50 | 32 | P | + | + | + | + | + | + | + | NA |
| **Boyacá** | Ventaquemada | B55 | 33 | P | + | + | + | + | - | + | - | MT738952 |
| **Cundinamarca** | Granada | C34 | 34 | NP | + | + | + | + | + | + | - | MT738953 |
| **Cundinamarca** | Granada | C23 | 35 | P | + | + | + | + | + | + | - | JX465125.1 |
| **Boyacá** | Arcabuco | B01 | 36 | NP | + | + | - | - | - | - | - | JX465121.1 |
| **Cundinamarca** | Silvania | MAP5 | | P | + | + | + | + | + | + | + | JX465121.1 |
| **Caldas** | Chinchina | Fol59 | | Tomato | + | - | - | - | - | - | - | MN745144 |
| **Uraba** | NA | FocR1 | | Banana | + | - | - | - | - | - | - | NA |
| **Guajira** | Dibulla | FocTR4 | |  | + | + | - | - | - | - | - | NA |

P= Pathogenic strain in cape gooseberry plants

NP= Non-pathogenic strain in cape gooseberry plants

NA= Non-available EF1a sequence
