## Supplementary Table 3 for "Novel effector genes revealed by the genomic analysis of the phytopathogenic fungus *Fusarium oxysporum* f. sp. *physali* (*Foph*) that infects cape gooseberry plants"

**Table S3**. Primer sequences used in this study

| **Primer Name** | **Sequence** | **Region amplified** | **Amplicon size bp** |
| --- | --- | --- | --- |
| SIX13-F | CGAATCCTTCATCATCGACA | Foph-SIX13 | 1142 |
| SIX13-R | AGGGTCTTGAGTTCGTTGACA |  |  |
| g7767.t1-F | atgaagcctgtggccattct | Eff1 | 493 |
| g7767.t1-R | ctaggttgcgaggatcacg |  |  |
| g10813.t1-F | atgaggttctccctatcacaga | Eff2 | 491 |
| g10813.t1-R | ctacttgaaacagccacctga |  |  |
| g11758.t1-F | atgcgttctaccgctatctctc | Eff3 | 519 |
| g11758.t1-R | taatcataaagccggatttcca |  |  |
| g11759.t1-F | atgaagtcaatcactctccttgc | Eff4 | 604 |
| g11759.t1-R | gtcggccaagtccgtcag |  |  |
| g14220.t1-F | ttcctgactctggccctgac | Eff5 | 343 |
| g14220.t1-R | ttagttgcagctccaatacgg |  |  |
| g14434.t1-F | atggttaaacacatccaactgc | Eff6 | 384 |
| g14434.t1-R | cccataataaccccctccaa |  |  |
| ￼g14470.t1-F | gaagttatcttacgccagcgttatt | Eff7 | 252 |
| g14470.t1-R | tcactagccacacaaacgatac |  |  |
