## Supplementary Figure 1 for "Novel effector genes revealed by the genomic analysis of the phytopathogenic fungus *Fusarium oxysporum* f. sp. *physali* (*Foph*) that infects cape gooseberry plants"

**A.**

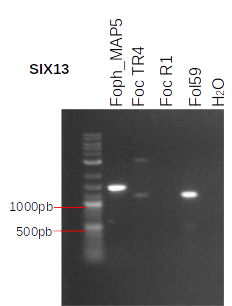

**B.**

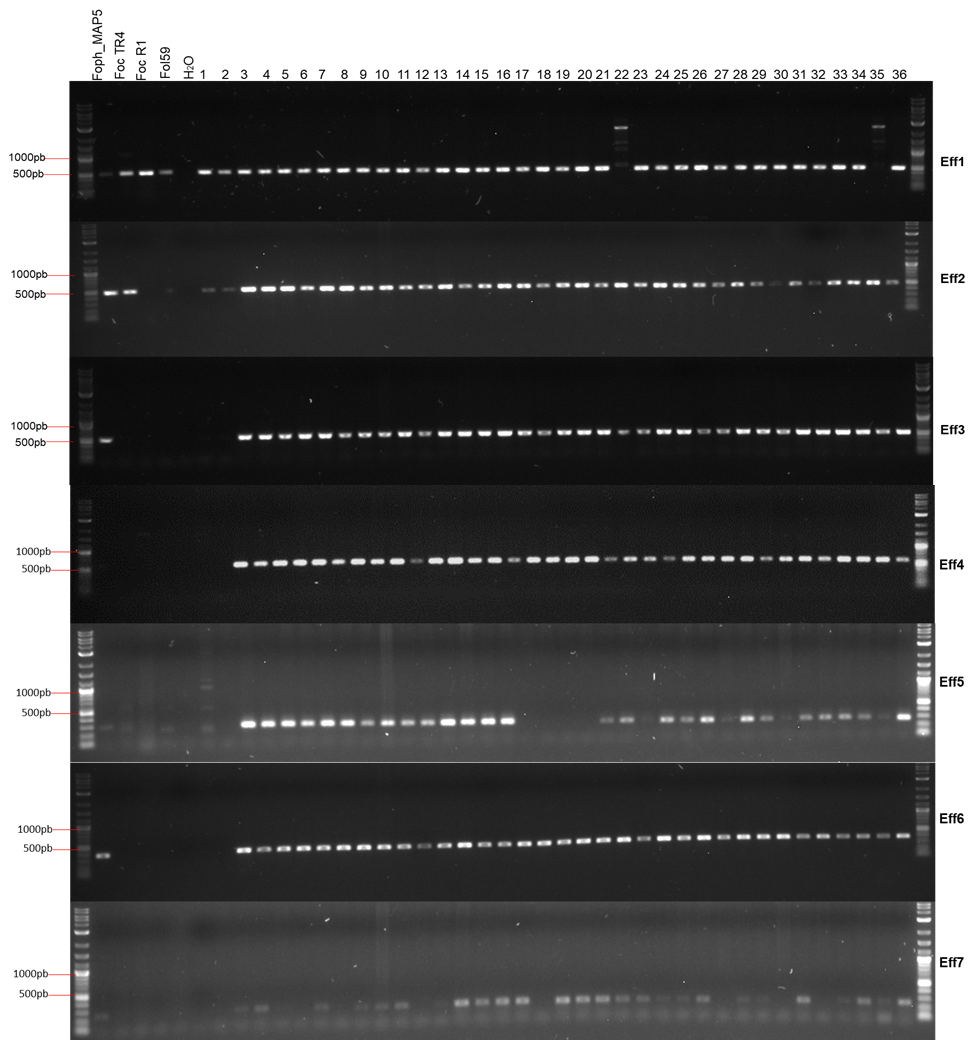

**C.**

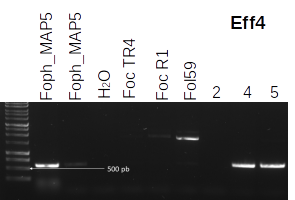

**Figure S1.** Confirmation of the novel *Foph* candidate effectors by PCR amplification **A.** Homologue of SIX13 **B.** Seven *Foph* novel candidate effectors (Eff1 to 7), showing their presence in a panel of 36 *F. oxysporum* isolates associated to the cape gooseberry crop (Table S2). **C.** Eff4 confirming its presence on *Foph*_MAP5. Control DNA: *Foph*_MAP5, Colombian isolates of Foc_TR4 (tropical race 4), Foc_R1 (race1) and Fol59. PCR product visualization was carried out following electrophoresis in a 1% agarose gel.
